# Comparison of wearable and clinical devices for acquisition of peripheral nervous system signals

**DOI:** 10.1101/2020.10.27.356980

**Authors:** Andrea Bizzego, Giulio Gabrieli, Cesare Furlanello, Gianluca Esposito

## Abstract

A key access point to the functioning of the Autonomic Nervous System is the investigation of peripheral signals. Wearable Devices (WDs) enable the acquisition and quantification of peripheral signals in a wide range of contexts, from personal uses to scientific research. WDs have lower costs and higher portability than medical-grade devices. But achievable data quality can be lower, subject to artifacts due to body movements and data losses. It is therefore crucial to evaluate the reliability and validity of WDs before their use in research. In this study we introduce a data analysis procedure for the assessment of WDs for multivariate physiological signals. The quality of cardiac and Electrodermal Activity signals is validated with a standard set of Signal Quality Indicators. The pipeline is available as a collection of open source Python scripts based on the pyphysio package. We apply the indicators for the analysis of signal quality on data simultaneously recorded from a clinical-grade device and two WDs. The dataset provides signals of 6 different physiological measures collected from 18 subjects with WDs. This study indicates the need of validating the use of WD in experimental settings for research and the importance of both technological and signal processing aspects to obtain reliable signals and reproducibility of results.

## 1 Introduction

The quantification of peripheral physiological nervous signals is a core step to measure the functioning of the Autonomic Nervous System [27]. Being able to observe these phenomena in real-life and without constraints imposed by laboratory settings is a key reason for adopting WDs in scientific research [19]. Wearable technologies enable the acquisition and quantification of physiological signals in a wide range of contexts, from personal uses to industrial and scientific research [46, 13, 33]. In a general sense, Wearable Devices (WDs) are portable, non-invasive devices that allow the acquisition of physiological signals during daily life, with no need of external equipment [6]. The application of WDs is boosted by their improved portability (e.g., increasing miniaturization and battery life) [14, 51], as well by new materials ([30, 11]) and layouts with reduced invasiveness, e.g.: epidermal tattoos [5], wireless ring pulse oximeter [22], garments [17] and masks [43].

Besides higher portability, WDs may have an apparent lower cost than clinical-grade devices, which enabled their diffusion for commercial applications [40]. In contrast to growing market availability, the adoption of WDs in research is still limited. Wearable technologies could open to the scientific community novel tools in the study of human physiology and autonomic responses [28, 27], paving the way for a new generation of experiments in which long-term monitoring and real-life ecological acquisitions are key aspects. In particular, a decrease in obtrusiveness on subject’s behaviour and daily activities is expected by adopting WDs instead of clinical devices [19]. Many scientific fields would benefit from the adoption of WDs for long-term, unobtrusive recording of physiological signals. In particular, scientific use of WDs is explored in affective computing [37, 35], research on autism [46, 26], interpersonal coupling [3, 8] and psychology in general.

However, due to the technical constraints imposed by improvements in miniaturization and autonomy of WDs, the data quality that can be achieved is typically lower and more subject to noise due to body movements [34, 50, 48]. Further, we need to consider the hurdles in extracting, preprocessing and analyzing the data from non-standard data interfaces: high scientific “Total Cost of Ownership”, possibly requiring manual preprocessing is also associated to limited reproducibility of research.

It is therefore necessary to create a working protocol and resources to validate WDs as a reliable scientific tool, before their use in research [23]. In particular, the validation should be based on open access scientific software and include a comparison with a baseline of results from signals collected by medical-grade devices, within the same or very similar experimental settings. In practice, a battery of signal processing algorithms or their composition in a pipeline should be applied to signals collected by clinical-grade and wearable devices, with a lower level of precision accepted considering the type and purpose of application.

Similar works in the literature mainly target particular settings; for instance, the usage of WDs in schools [42], with a particular type of users [24]. Studies focusing on the comparison between clinical and WDs are usually application-specific; for instance, to study physiological synchrony [44], sleep [16, 15], vital signs monitoring [12] or usage in clinical settings [4, 2]

However, despite its importance, the validation of a WD is often omitted or left to manufacturers, who rarely provide the dataset used and the details about the validation procedure. Although several datasets with physiological signals are available, such as DEAP [25], MANHOB-HCI [39], SEMAINE [32] and the PhysioBank archive [21], none of the existing resources provides signals collected from both wearable and clinical grade devices to allow the comparison.

Finally the set of algorithms applied in the validation should be also available for reproducibility and further reference.

In this study we introduce scientific software and data aimed at validating the use of WD for research within different real-life contexts (resting and movement), focusing on cardiac and electrodermal activity signals. Our contributions are: (a) creation of the Wearable and Clinical Signals (WCS) dataset which allows comparing a representative set of WD with clinical-grade devices; (b) a procedure to assess the validity of signals collected from WD; and (c) quantitative results on physiological signals of interest for the affective-computing field.

Physiological signals were collected from a clinical-grade device and from two WDs to allow both the direct comparison of the signals’ data quality and the investigation of the reproducibility. The adopted protocol comprises two different activities to evaluate the performance in different real-life contexts.

## 2 Methods

### 2.1 Participants

A total of 18 participants (6 Males) were recruited for the experiment through announcements among university students (Age: Average=27.6, SD=7.04). The participation to the experiment was voluntary and granted credits for the fulfillment of an academic course. Before starting the experiment each participant was briefed about the purpose and procedure of the experiment and gave the informed consent. The experiment was approved by the Independent Review Board of the University of Trento (2017-019) and was conducted according to the principles of the Declaration of Helsinki. All the data have been de-identified by assignment of randomized numeric identifier to each subject and we removed any reference to the absolute timestamps.

### 2.2 Devices and architecture

The physiological signals were simultaneously collected by three devices (see Figure 1): the FlexComp unit which provided the reference signals, and two WDs: the Empatica E4 and the ComfTech HeartBand.

**Figure 1:**
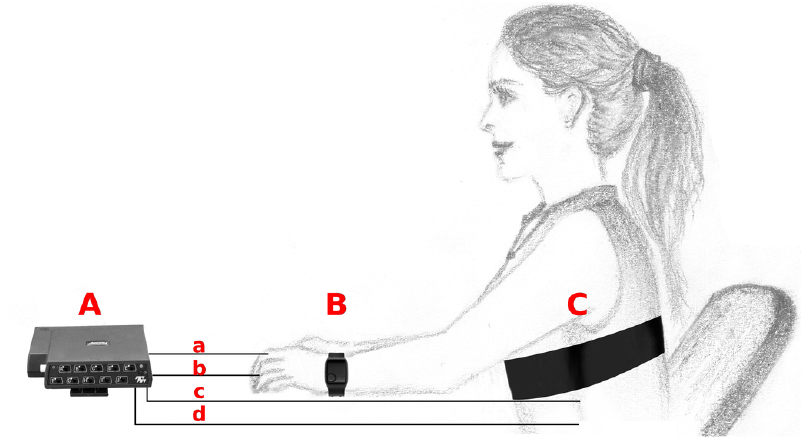
Illustration showing the devices used for the acquisition of physiological signals. A: Thought Technology FlexComp (picture from www.thoughttechnology.com) with four wired sensors: a) Electrodermal activity, b) Blood Volume Pulse, c) Electrocardiogram, d) Respiration; B: Empatica E4 (picture from www.empatica.com); C: ComfTech HeartBand.

The FlexComp acquisition unit was connected through optical fiber to a signal converter which in turn was connected to a Windows 10 workstation where the proprietary BioGraph Infinity Software Platform managed the acquisition and collection of the signals from the unit.

The two WDs sent the data through Bluetooth connection to an Android (5.1) tablet (Samsung Tab A) where a specifically developed app gathered the signals and sent them to a database through secure connection. The app interface also allows the annotation of manual markers to allow the synchronization of the signals collected with the FlexComp and WDs.

Before starting the experiment, the signals acquired through the two WDs were visually inspected by activating the real-time plotting function.

The WDs used in the experiment were selected according to four requirements we defined to explicitly highlight the key characteristics that allow the usage in everyday life:

1. Wearability: the device should be easily worn by the subject with no need of applying conductive gels, wired electrodes or similar preparations;
2. Streaming: the device should stream the data in real-time to an external collector through Bluetooth connection;
3. Availability: the device should be commercially available at the moment of testing (prototypes, proof-of-concepts or custom devices were not considered);
4. Performance: the final choice should favor the device with higher sampling rate and higher number of sensors to select the most advanced technological solution.

Wearability is the main characteristics of WDs: it facilitates the acquisition of signals, reducing the discomfort of users and the time required to set the experiment, and it allows expanding the investigation to everyday applications. In some experimental settings, WDs are expected to be set directly by the user, with no aids or supervision from an experimenter. For this reason, the WDs to be selected should be easily worn by the user, without requiring any special preparation of the device. Streaming capability is also a key feature of WDs: it allows synchronization and communication with other devices and opens to real-time applications which are fundamental in remote health monitoring. In the experimental settings of this study, this requirement was also imposed by the specific sensing architecture, with one hub collecting the data from all wearable devices and synchronizing them with the data from the FlexComp unit. There are several types of WDs currently available: from prototypes to medical-grade devices. Prototypes are devices created to demonstrate a novel technological solution or developed for a specific purpose or application. While some of these prototypes might be made commercially available through start-ups or small companies, their usage requires some technological skills and, for this reason, fail to comply with the “wearability” requirement. On the opposite side, medical-grade WDs are the most advanced solution, but were not considered in this study as these types of devices already undergo a set of validation studies to be certified as “medical-grade”. We focused instead on intermediate solutions that should be commercially available and could be accessed even by researchers with limited technical skills. Finally, we selected those devices that, compared to similar solutions, offered the best performances in terms of signal acquisition. We considered the maximum sampling frequency and the number of different signals that could be collected by the same device.

After a preliminary phase in which a list of WDs was tested, we selected two devices, that full-filled all the requirements: the Empatica E4 and the Comftech HeartBand. Empatica E4 was chosen as a representative of wrist-band devices, while Comftech HeartBand belongs to the smart-garments category.

#### 2.2.1 Reference: Thought Technology FlexComp

The clinical device we used as reference was the Thought Technology FlexComp unit (commercial code: T7555M). It is a customizable acquisition unit which provides up to 10 input slots that can be used to connect diverse physiological sensors. Maximal sampling frequency is 2048 Hz, which was adopted for the experiment, with 14 bits of resolution for each input. The physiological signals acquired with the FlexComp unit were:

1. Electrocardiogram (ECG): using three electrodes placed over the left and right coracoid processes and below the ribs on the left. The signal is pre-amplified and filtered by the EKG Sensor (T9306M) which returns a single channel read in millivolts. UniGel electrodes (T3425) are used as conductive mean between the sensor and the skin;
2. Electrodermal Activity (EDA): two finger bands with Ag-AgCl electrodes (SA2659) are placed on the second and fourth finger of the left hand and connected to the sensor (SA9309M). The skin conductance is measured in microSiemens (*µ*S);
3. Blood Volume Pulse (BVP): the sensor (SA9308M) is placed on the third finger of the left hand. The relative amount of reflected infrared light is measured;
4. Respiration (RESP): a band is worn on the chest to measure the relative volumetric expansion by elongation of an elastic patch (SA9311M);
5. Trigger (TRG): a handle with a button to generate electrical impulses used to manually mark the experimental events.

The signals collected from the FlexComp unit were the raw signals provided by the device and no signal pre-processing was applied, except for the conversion to the signal units.

#### 2.2.2 Empatica E4

The Empatica E4 [20] is a multi-sensor wristband designed for real-life acquisitions of physiological signals. It has streaming (through Bluetooth Low Energy) and storing (internal flash memory) capacity; communication with an Android smartphone is provided by the EmpaLink Software Development Kit which is used by a customized app for signal acquisition. The collected signals are:

1. BVP: four Light Emitting Diodes (LEDs) are used to generate light at two different wavelengths (Green and Red) and two photodiodes are used to measure reflected light. Using two wavelengths and an appropriate proprietary algorithm to preprocess the signals should reduce motion effects and sensitivity to external sources of light. Sensors are placed on the bottom of the wristband case in firm contact to the skin; signal is sampled at 64 Hz;
2. EDA: two stainless steel electrodes are placed on the band to allow positioning on the inner side of the wrist. Skin conductance is measured in microSiemens at 4 Hz sampling rate;
3. Acceleration (ACC): three axes acceleration (range ± 2g) is measured at 32 Hz sampling frequency;
4. Skin Temperature (ST): measured by an infrared thermopile placed on the back of the case, at 4 Hz sampling frequency.

The collected signals were the raw signals provided by the device and no signal pre-processing was applied, except for the conversion to the signal units.

#### 2.2.3 ComfTech HeartBand

The Comftech HeartBand is an elastic band with embedded tissue electrodes and an acquisition unit that is connected to the band through two snaps. Streaming is based on Bluetooth 2.0 and the documentation to decode the hexadecimal messages from the device was provided by the manufacturer.

The collected signals are:

1. ECG: using two tissue electrodes placed over the chest and connected to the acquisition unit where the signal is sampled (128 Hz), pre-amplified and filtered to return a single channel read;
2. ACC: three axes acceleration is measured by a sensor embedded on the acquisition unit; sampled at 200 Hz;

The collected signals were the raw signals provided by the device and no signal pre-processing was applied, except for the conversion to the signal units.

### 2.3 Experimental protocol and procedure

After the signed informed consent was given, the experimenter instructed the participant about the placement of WDs and the experimental settings. The setup of the FlexComp unit, of the acquisition software and the positioning of the sensors was performed by the experimenter to ensure high quality signals. Regarding the WDs, the participant was asked to place himself the devices, after having received detailed instructions from the experimenter. This procedure reflects the idea that WDs are to be used in real-life context, where the presence of an experimenter is not expected. In particular, the participant was allowed to adjust the positioning of both the Empatica E4 and of the ComfTech HeartBand to favor the comfort during the experiment. However, the placement of the E4 wristband was inspected by the experimenter to ensure the device was not worn too tight (which would have prevented the physiological blood flow) or too loose (which would have prevented the smartwatch case to be in contact with the skin) to guarantee optimal conditions for a correct BVP signal acquisition.

The experiment comprised two phases, baseline (300 s) and movement (300 s), which were administered following the same order for all participants. During baseline the participants were asked to remain sit and still. The signals collected during this phase represent a reference for the maximal signal quality that can be achieved by WDs, as all external sources of noise that could affect the signal quality (such as moving artifacts, sensor displacement or detachment) are experimentally avoided. This condition is far from the real-life context in which the WDs are expected to be used, but it is needed to isolate the effects of technical limitations and constraints from other causes of errors.

The effects of body movements, instead, can be better evaluated on the movement phase which concludes the experiment. During this phase the subject is asked to stand and simulate a walking on place, as the free movement was prevented by the wires connecting the medical-grade sensors with the FlexComp unit. The experimenter suggested moving naturally to replicate the intensity of usual walking. In particular, special attention was paid to movements of the left hand as, due to presence of sensors, participants tended to keep it still and near to the body. This experimental phase is expected to be more similar to real-life contexts and can be used to evaluate performances of algorithms for every-day applications, such as health monitoring, fitness and stress detection.

### 2.4 Preprocessing

The raw signals have been preprocessed in order to appropriately format the data for the distribution and signal analysis. The first step was the synchronization of the two signal flows, from the FlexComp device and from the WDs. After the synchronization the absolute timestamps of the samples were converted into elapsed time since the start of the experiment (start of the baseline). Based on the TRG signal the instants of the beginning and end of each experimental phase were extracted and used to split the signals. Then each portion of signal was separately saved. The acceleration signals, which are three-dimensional, were converted into uni-dimensional signals by computation of the acceleration module.

Due to the sensitive nature of the data, the dataset is not public, but the collected signals are available to researchers aiming to create and test new algorithms for the automatic identification of noise and artifacts: https://doi.org/10.21979/N9/42BBFA.

### 2.5 Signal Quality analysis

Assessing the quality of the collected signals is crucial for reproducible research; this step is even more important when working with WDs, as signals are more sensitive to artifacts and noise due to body movements and technical constraints (see Figure 2).

**Figure 2:**
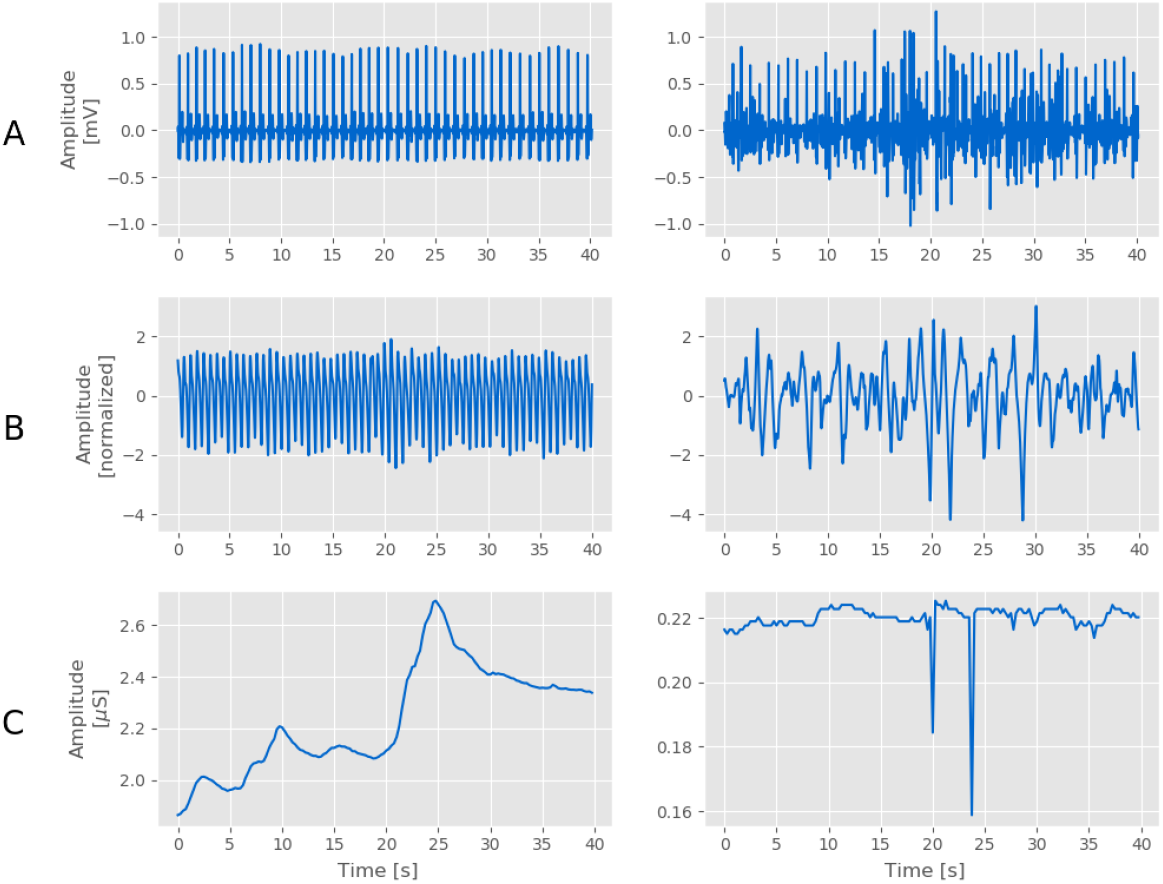
Portions of signals from WDs extracted from the WCS dataset to show examples of good (left column) and bad (right column) quality signals. A: ECG signals collected by the HeartBand; B) BVP signals collected by the E4; C) EDA signals collected by the E4.

In this study, we adopted a data analysis pipeline (Figure 3) to investigate the quality of the acquired cardiac and EDA signals and of derived signals and indicators. A set of Signal Quality Indicators (SQIs) has been defined to compare the quality of signals from clinical-grade and wearable devices. The SQIs have been computed on all cardiac signals and EDA signals of the baseline and movement sessions. A Generalized Linear Model (GLM) was fit to investigate the contribution of the type of device (clinical or wearable) and session (baseline or movement) and their interaction on each SQI.

**Figure 3:**
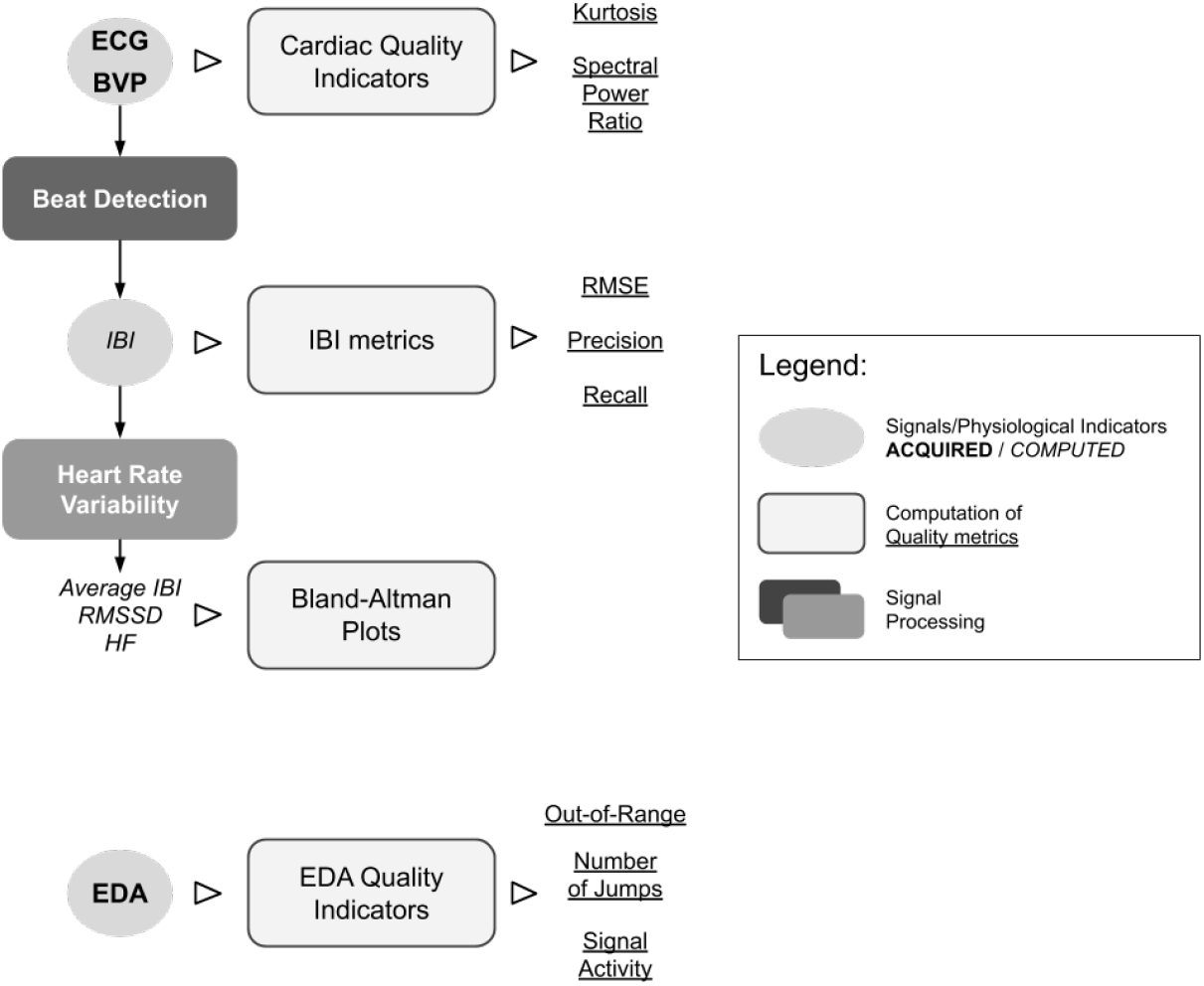
Data Analysis procedure adopted in this study.

The processing pipeline was developed in Python, based on pyphysio [9], and it is publicly available at https://gitlab.com/abp-san-public/wearable-clinical-devices.

#### 2.5.1 Cardiac signals

Several Signal Quality Indicators (SQIs) have been proposed in literature to evaluate the quality of cardiac signals. The SQIs can be grouped in two main categories: indicators that require a previous detection of the heart beats, and indicators that are computed directly on the signal. We first used two SQIs from the second category to obtain an overall quantification of the signal quality, then adopt a specialized beat detection algorithm to assess the possibility of effectively identify the heart beats.

We used the Kurtosis (*K*) and the Spectral Power Ratio (*ψ*) which have been proposed for both the ECG [29, 7] and the BVP [18, 47]. Following the thresholds proposed in the literature, good quality signals are expected to have *K >* 5 in case of ECG [29], while *K <* 3.5 in case of BVP [38].

The Spectral Power Ratio (SPR) *ψ* is defined as 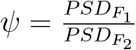 where *PSDF* is the spectral power in the frequency band *F*. *F*_1_ is associated with the components in the signals that convey the information about the heart beats (the QRS complexes in the ECG and the pulses in the BVP); *F*_2_ is the band which contains all the main frequency components of the signals. For ECG signals: *F*_1_ = [5, 14]*Hz, F*_2_ = [5, 50]*Hz*, [29]; for BVP signals: (*F*_1_ = [1, 2.25]*Hz, F*_2_ = [0, 8]*Hz* [18]). *ψ* is expected to have values 0.5 *≤ ψ ≤* 0.8 [29].

To compute the Signal Quality Indicators (SQIs) the signal is resampled (128 Hz) and filtered (band pass filter: [0.5, 50] Hz). Then the signal is standardized and segmented with 5 s length non-overlapping windows. The reported *K* and *ψ* are the average value across all segments.

We then investigated the reliability of the acquired cardiac signals in measuring the cardiac activity. To this aim we applied a beat detection algorithm to identify the heart beats and derive the Inter Beat Intervals (IBIs) series, which is the series of distances between consecutive beats. The validation metrics are then computed by comparing the IBIs series with the reference IBIs series obtained from the ECG signal of the Flex Comp, which were manually validated. These metrics are different from the SQIs presented before, as they are computed on a derived signal and not on the original cardiac signal, and thus the results depend also on the performance of the beat detection algorithm.

The ECG signals from the Flex Comp unit were used to obtain the reference IBIs series used for the validation: heart beats were detected with the Adaptive Beat Detection (ABD) algorithm (see 2.5.2) and then visually inspected to manually correct false and misdetections.

The ABD algorithm was applied also to the ECG signals of the Heart Band, while the Derivative-Based beat detection algorithm [10] was used to derive the IBIs series from the BVP signals of both the Flex Comp and E4. All IBIs series were then processed to find detection errors by application of the Adaptive Outlier Detection (AOD) algorithm (see 2.5.3). In summary, for each subject we obtained the reference IBIs series (*IBI*_*r*_) and those derived from the ECG signal of the Heart Band (*IBI*_*HB*_), from the BVP signal of the Flex Comp (*IBI*_*F C*_) and from the BVP signal of the E4 (*IBI*_*E*4_).

The quantification of the reliability of the beat detection was assessed in terms of precision (*P*) and recall (*R*) of the detected beats and by root mean square error (*e*_*RMSE*_), by comparing each derived IBIs series (*IBI*_*d*_) with the *IBI*_*r*_. To compute these metrics, we paired each beat *b*_*d*_ of the *IBI*_*d*_ to a beat *b*_*r*_ of *IBI*_*r*_. The pairing is considered valid if |*b*_*d*_ *− b*_*r*_ | *<* 0.5 s. The beats that were successfully paired are considered True Positives (TPs), the remaining unpaired beats in *IBI*_*d*_ are considered False Positives (FPs) and the remaining unpaired beats in *IBI*_*r*_ are considered False Negatives (FNs). Then we counted the number of TPs (*n*_*TP*_), the number of FPs (*n*_*FP*_) and the number of FNs ((*n*_*FN*_)) to compute the overall precision *P* = *n*_*TP*_*/*(*n*_*TP*_ + *n*_*FP*_) and recall *R* = *n*_*T P*_ */*(*n*_*TP*_ + *n*_*FN*_). The RMSE is computed on paired beats: 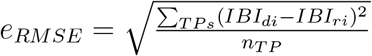.

Finally, we evaluated the impact of the different types of WDs and cardiac signals on the main physiological indicators used to quantify the Heart Rate Variability (HRV). For each IBI signal we computed: (a) the average of the IBI, (b) the Root of the Mean of the Squares of the Subsequent Differences (RMSSD) between the IBI, and (c) the relative power in the High Frequency (HF) band (between 0.15 and 0.4 Hz) [31]. The three HRV indicators are computed for each subject and session, on all cardiac signals. The HRV indicators computed from the IBI extracted from the ECG signals collected with the FlexComp unit are used as reference. The HRV indicators computed from the IBI extracted from all other cardiac signals are qualitatively compared to the reference HRV indicators using Bland-Altman plots. Differences are reported in terms of percentage to allow comparing the performances with the different HRV indicators.

#### 2.5.2 Adaptive Beat Detection

The Adaptive Beat Detection (ABD) algorithm applied to the ECG signal is composed of two steps:

i. Computation of the local range of the signal;
ii. Peak detection with threshold from the local range.

In the first step, the signal is segmented by windowing (windows width: 1 s, overlap: 0.5 s) and the range is computed for each segment. The result is the estimation of the local range of the signal.

In the second step, the signal is scan to find local maxima; a local maximum is considered a heart beat (i.e.: R peak of the QRS complex) if it is followed by a local minimum and the difference is greater than 0.7 times the local range of the signal.

#### 2.5.3 Adaptive Outlier Detection

The AOD uses a fixed size cache vector *IBI*_*c*_ = (*ibi*_1_, *ibi*_2_, …, *ibi*_*k*_) to store the last *k* valid IBI values, in order to adapt to IBI variability. The size *k* is empirically set to 5 and the cache is initialized with the median value of the IBIs series. Outliers detection is also regulated by the sensitivity parameter *ϕ*, which is empirically set to 0.25.

A detected beat is considered valid if its corresponding IBI value is within the interval [(1-*φ*)*ibi*_*median*_, (1+*φ*)*ibi*_*median*_], where *ibi*_*median*_ is the median of the values in *IBI*_*c*_. When a new valid beat is detected, its IBI value is used to update *IBI*_*c*_ using the First-In-First-Out rule. *IBI*_*c*_ is re-initialized when *k* consecutive non valid beats are detected.

#### 2.5.4 Electrodermal Activity signals

Only few studies address the problem of assessing the quality of EDA signals as this is a quite recent issue, associated with the emergence of wearable devices for EDA acquisition. The main approaches are based on machine learning to classify good and bad quality portions of the EDA starting from a list of computed metrics [41, 49], but without identifying the most appropriate that, alone, could be used as SQI. On the other hand, these models are not very useful with signals from new devices as they need to be re-trained on new data that would require a manual labelling process.

#### Quality metrics of EDA signals

In this study, we adopted two metrics proposed by Kleckner and colleagues [23]: the Ratio of Out-of-Range samples (*r*_*oor*_) and the number of jumps, or drops, in the signal (*n*_*j*_).

Following [23], we define *r*_*oor*_ as the ratio between the number of samples with value exceeding 60 *µS* or below 0.05 *µS* and the total number of samples. We empirically defined the threshold *r*_*oor*_ = 0.05 to identify good (*r*_*oor*_*leq*0.05) and bad (*r*_*oor*_ *>* 0.05) signals; the selected threshold corresponds to a percentage of 5% of acquired samples that exceed the physiological range for EDA signals.

A *jump* is defined in [23] as a portion of the signal in which the absolute value of the derivative of the signal is grater than 10 *µS/s*. This threshold was defined on signals sampled at 32 Hz, but the EDA signal from the Empatica is acquired at 4 Hz which make the 10 *µS/s* threshold too high to allow the correct identification of the jumps. In addition, the amplitude of the EDA signal, and thus its derivative, also depends on the characteristics of the device, such as: amplification, type of electrodes and positioning. For these reasons, we derived a normalized version of the metric proposed by Kleckner and colleagues.

We first applied a low pass filter to the EDA signal (cutoff frequency 0.05 Hz) to remove the trend and the long-term components, obtaining the filtered signal (*s*_*f*_); then computed the normalized derivative 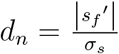, where *s*′_*f*_ is the derivative of the filtered signal and *σ*_*s*_ is the standard deviation. Then we considered as jumps the samples where *d*_*n*_ is greater than 10. We considered as acceptable a total number of 5 jumps over the 5 minutes sessions of the baseline and movement: therefore signals with *n*_*j*_ *≤* 5 (corresponding to maximum a jump per minute of acquisition) are considered with good quality. In addition, we defined a new metric, called Signal Activity metric (*α*), to account for the cases in which the amplitude of the peaks associated to the Galvanic Skin Response is too low or the peaks are not present (see Figure 2C). We defined 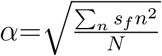 where *s*_*f*_ is the filtered signal and *N* is the number of samples. As a threshold value we selected *α* = 0.05 *µS* which is the minimal amplitude for a peak in the EDA signal to be considered as part of the Galvanic Skin Response, as proposed in [1]. EDA signals with *α <* 0.05 *µS* are considered of bad quality.

Before computing the SQIs, the EDA signals are filtered (low-pass filter, cutoff frequency *f*_*lp*_ = 1.5 Hz). The EDA signals from the Flex Comp device are first downsampled to 4 Hz to allow the comparison with the signals from the Empatica E4. The SQIs are computed on segments of the signals obtained from non-overlapping 20 *s* windows. The reported *r*_*oor*_ and *α* are the average of the values computed for each segment; the *n*_*j*_ is the sum of the jumps in each segment.

## 3 Results

### 3.1 Cardiac signals

In general, the ECG signals collected with both the Flex Comp and the Heart Band (Figure 4A) showed good quality in terms of Kurtosis although body movements cause a degradation; ECG signals from the HeartBand seems to contain spurious frequency components as indicated by the higher SPR. The BVP signals from both the Flex Comp and the E4 (Figure 4B) showed a very good Kurtosis; however the low SPR for the movement session indicates that this signal is highly affected by body movements, as already reported in literature [48].

**Figure 4:**
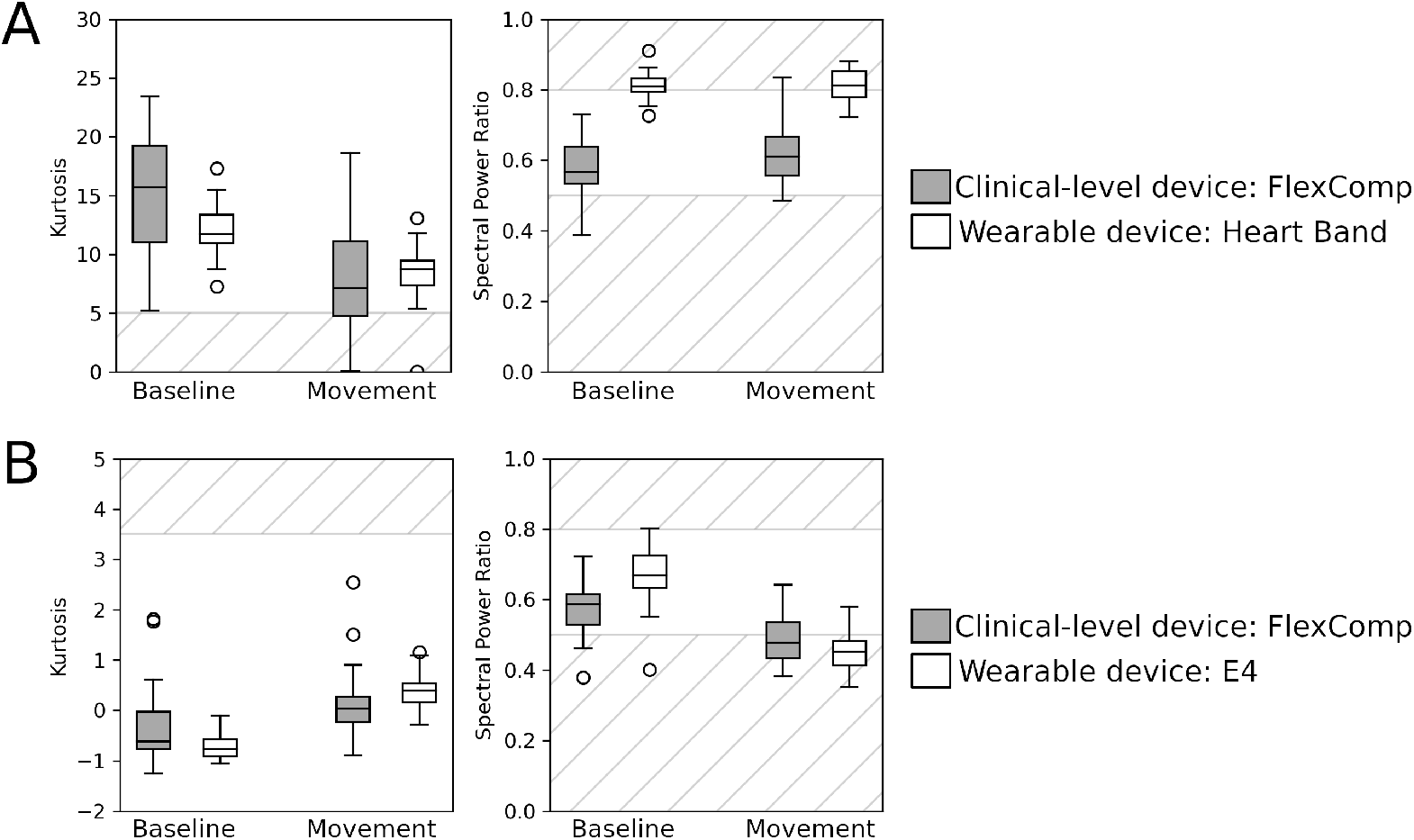
SQIs of the cardiac signals: A. Kurtosis and Spectral Power Ratio of the ECG collected from the FlexComp (gray) and HeartBand (white); B. Kurtosis and Spectral Power Ratio of the BVP collected from the Flex-Comp (gray) and E4 (white). Striped areas indicate the ranges of SQI values associated with low signal quality.

For the ECG signals, Kurtosis was found to be significantly affected by the type of session (*p <* .001) and type of the device (*p* = 0.035), but the latter does not survive the Bonferroni correction for multiple tests. A significant contribution of the type of device was found on the SPR (*p <* .001). For the BVP signals, a significant contribution of the type of device (*p* = .038) and of the interaction between type of device and session (*p* = .020) was found on the Kurtosis; both results would not survive Bonferroni correction. On the SPR, a significant contribution was found for the type of device (*p <* .001), type of session (*p* = .001) and their interaction (*p <* .001).

During the baseline sessions all devices show good performances in terms of *P* and *R*, although the recall of the IBIs series derived from the BVP signals is inferior (see Figures 5. The *e*_*RMSE*_ is also sensibly higher for the IBIs series derived from the BVP signals. This result is expected as the beat detection in BVP signals is in general more difficult due to posture and individual physiological characteristics [36]. As expected, all metrics worsen during the movement sessions, in particular the IBIs series derived from BVP signals of the Empatica E4 show very low recall (below 0.4) and high *e*_*RMSE*_.

When considering the Bland-Altman plots of the physiological indicators extracted from the cardiac signals (Figure 6), it emerges that BVP signals show the greatest differences, with the E4 performing worse than the clinical level device.

**Figure 5:**
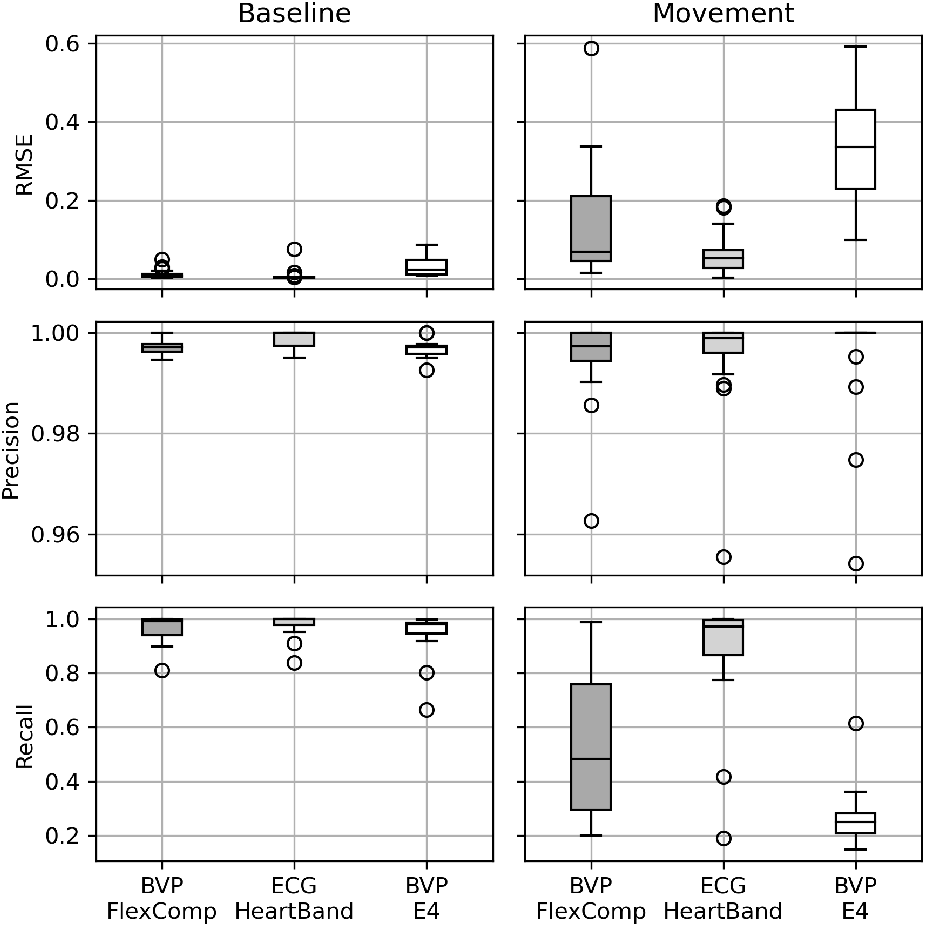
Metrics of the quality of the Inter Beat Intervals extracted from the cardiac signals.

**Figure 6:**
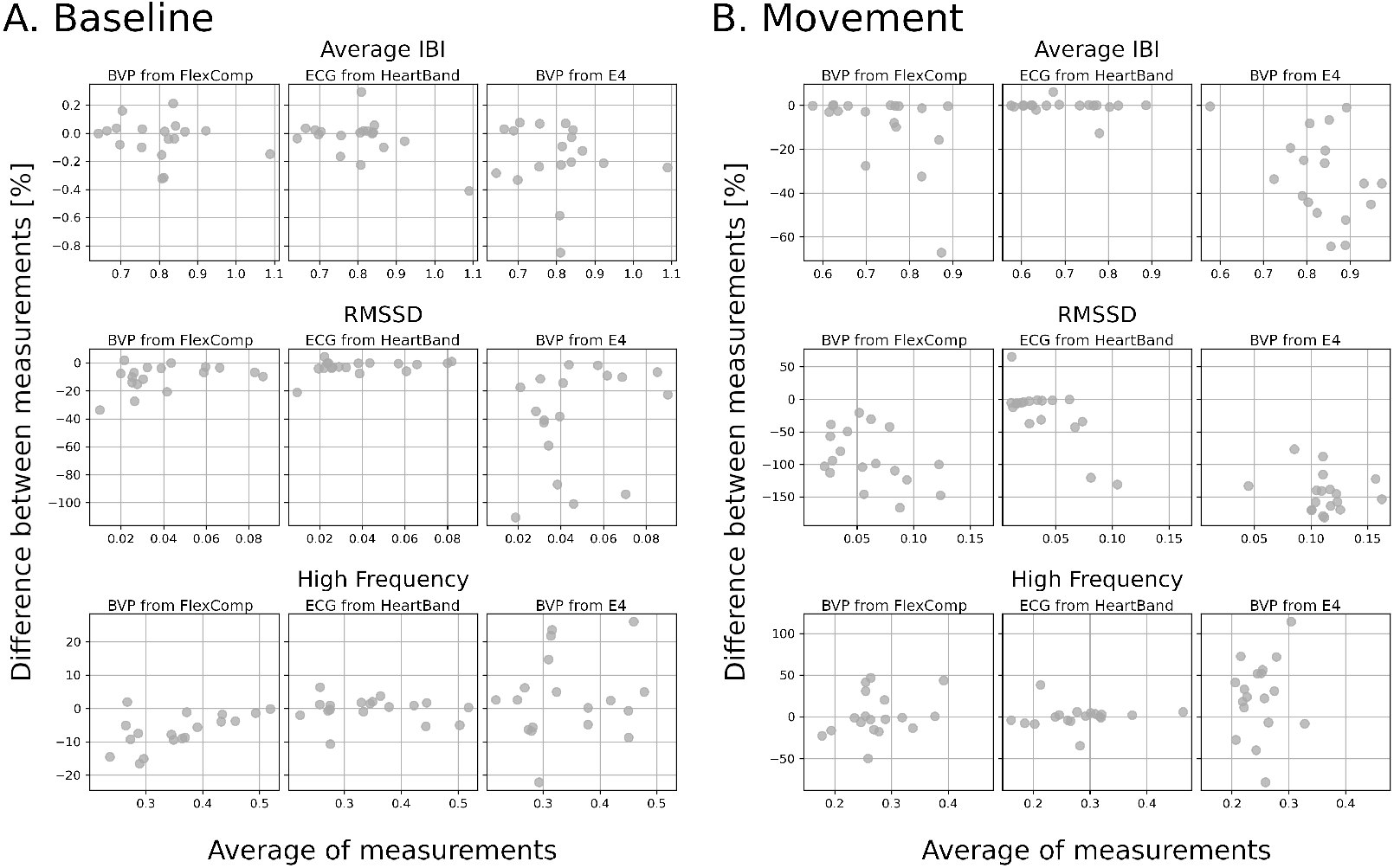
Bland-Altman plots to assess the reliability of the physiological indicators computed from the IBI signals extracted from different signals and devices. All plots refer to the indicators computed from the ECG signals collected by the FlexComp.

### 3.2 Electrodermal activity

When we consider the SQI of the EDA signals (Figure 7), we observe that none of the EDA signals collected with the Flex Comp presented Out-of-Range samples while few signals collected by the Empatica E4 appear corrupted, with a worsening effect due to the body movements. In terms of number of jumps, the Flex Comp unit provided good quality signals for both the baseline and the movement sessions, with few exceptions. Instead, the signals collected with the Empatica E4 presented a low quality with an average *n*_*j*_ = 31.6 for the baseline and *n*_*j*_ = 36.7 for the movement session: more than one jump every 10 s. The Signal Activity index of the signals collected with the Empatica E4 is low, despite the high number of jumps, in particular for the baseline session.

**Figure 7:**
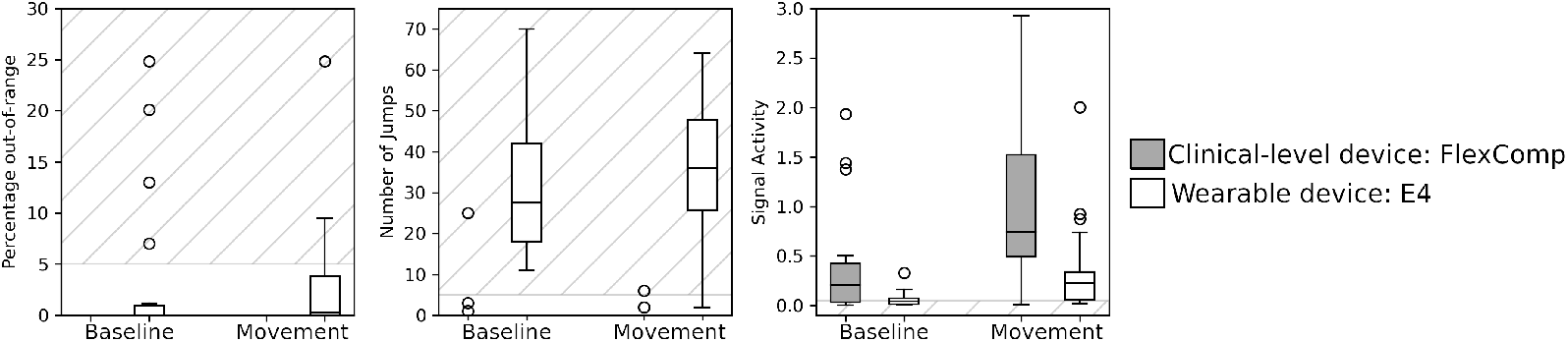
SQIs of the EDA signals collected from the FlexComp (gray) and E4 (white). Striped areas indicate the ranges of SQI values associated with low signal quality.

The GLM revealed a significant contribution of the type of the device on the Percentage of Out-of-Range samples (*p* = .024, which would not survive Bonferroni correction), and on the Number of Jumps (*p <* .001). Only the Signal Activity showed a main effect of the type of session (*p <* .001).

## 4 Discussion

We adopted a set of SQIs derived from the literature to assess the overall quality of the cardiac signals and EDA signals provided in the WCS dataset. In general, the signals collected with the clinical device showed an optimal signal quality on the baseline session, indicating that the experimental design and settings were appropriate for the acquisition of scientific-level data. During the movement session, the induced artifacts slightly worsen the signal quality, in particular for the BVP signal, which is known to be more sensitive to body movements. During the baseline, the wearable devices collect cardiac signals with a quality comparable to the signals from the FlexComp unit but are more affected by the body movements as shown by the worse signal quality of the movement session. For ECG signals, the Kurtosis is more sensitive to decrease in signal quality due to movements, while the SPR is capable of sensing differences between clinical and WDs. The SPR of BVP signals is influenced by both differences between the type of session (baseline and movement) and of devices (WDs and clinical); Kurtosis was found to be less effective in quantifying differences between devices and sessions.

These patterns are confirmed when analysing the precision of physiological indicators through Bland-Altman plots: the highest precision, during both baseline and movement precision, is achieved with the ECG signal from the HeartBand, while for the indicators computed from BVP signals the precision is lower, for both the FlexComp unit and E4. The signal quality of the EDA signals collected by the E4 do not appear to be appropriate for scientific research on our data: all SQIs detected significant differences between the quality of signals collected with the FlexComp unit and the E4; only the Signal Activity indicator resulted sensitive to differences between baseline and movement.

The lower quality of signals collected from WDs can be attributed to a number of technical limitations. For instance, the performance of some sensor components are reduced to decrease battery consumption and increase autonomy. In particular, the sampling frequency is kept lower than medical-grade devices, usually below 256 Hz with consequent issues in the accuracy of the physiological indicators computed. To favour wearability, no aids are used to fix the sensors to the body (e.g. adhesive ECG electrodes). This allows for minimal shifts of the sensors that can introduce noise on the acquired signal [34, 50, 48]. Finally, although some anatomical loci are more appropriate than others for measuring a physiological signal, the need of embedding the sensor in a wearable and comfortable support might constrain the positioning of the sensor into a non-optimal locus where the signal magnitude might be lower, such as the wrist instead of hand palm in the case of EDA [45].

This study evidenced the need of testing and validating the use of WD in experimental settings for research and the importance of both technological and signal processing aspects to obtain reliable signals. The validation should reflect the experimental context in which the data are collected; in particular, since the WD are employed to favor the freedom of movement of the subject, the influence of artifacts due to movements should be carefully considered.

The use of proper SQI represents a quantifiable and reproducible method to identify the signals with good quality. However, the evaluation of the applicability should be done for each use case, to account for the specific aim of the investigation and requirements in terms of reliability of the computed metrics. The analysis of the Bland-Altman plots shows that different signals and devices are capable of different levels of precision of the computed physiological indicators. The scope of the analysis should be considered when assessing whether the precision required is compatible with the confidence ranges.

The collected data and the code to reproduce the pipeline are offered to the scientific community to encourage the development and validation of new SQI and algorithms. The adoption of standardized metrics and procedures should favor the implementation of experiments with multivariate physiological signals, based on WDs.

As a final note, it is important to remark that the results of this study are not to be interpreted in terms of technological validation of the WDs used for the experiment. The technological development is still in progress and manufactures of WDs are constantly improving the sensing and acquisition technologies. Therefore results reflect the state of the technology at the time of the experiments. Nevertheless, the validation and quantification of the quality of the collected data is a critical step that should be performed each time the technology or the WDs are updated.

## Author contribution

Conceptualization, A.B., C.F. and G.E.; methodology, A.B.; software, A.B. and G.G.; resources, G.E.; data curation, G.G.; writing–original draft preparation, A.B.; writing–review and editing, A.B. and C.F. and G.E.

